# Patterns of gene flow across multiple anthropogenic infrastructures: Insights from a multi-species approach

**DOI:** 10.1101/2019.12.16.877670

**Authors:** Jonathan Remon, Sylvain Moulherat, Jérémie H. Cornuau, Lucie Gendron, Murielle Richard, Michel Baguette, Jérôme G. Prunier

## Abstract

Large-scale Transportation Infrastructures (LTIs) are among the main determinants of landscape fragmentation, with strong impacts on animal dispersal movements and metapopulation functioning. Although the detection of LTIs impacts is now facilitated by landscape genetic tools, studies are often conducted on a single species, although different species might react differently to the same obstacle. Multi-specific approaches are thus required to get a better overview of the impacts of human-induced fragmentation. We surveyed four species (a snake, an amphibian, a butterfly and a ground-beetle) in a landscape fragmented by six LTIs: a motorway, a railway, a country road, a gas pipeline, a power line and a secondary road network. We showed that half of the overall explained genetic variability across all species was due to LTIs. While the butterfly was seemingly not impacted by any LTI, the genetic structure of the three other species was mostly influenced by roads, motorway and railway. The power line did not affect any species and the gas pipeline only impacted gene flow in the ground-beetle through forest fragmentation, but other LTIs systematically affected at least two species. LTIs mostly acted as barriers but we showed that some LTIs could somehow promote gene flow, embankments probably providing favourable habitats for vertebrate species. Considering the high variability in species response to LTIs, we argue that drawing general conclusions on landscape connectivity from the study of a single species may lead to counterproductive mitigation measures and that multi-species approaches should be more systematically considered in conservation planning.

## 2. INTRODUCTION

The human-induced fragmentation of natural habitats is one of the main determinants of the global biodiversity collapse (Fahrig, 2003). The most ubiquitous form of habitat fragmentation is due to large-scale transportation infrastructures (LTIs; Forman & Alexander, 1998). LTIs are linear infrastructures allowing the transportation of goods, vehicles or energy, such as roads, motorways, railways, power lines, pipelines and canals. They are expanding considerably, creating dense transportation networks with profound impacts on natural ecosystems. It notably deeply affects metapopulation dynamics through a reduction in population sizes in response to direct habitat degradation but also through a reduction in demographic and genetic exchanges between populations in response to a decrease in the permeability of the landscape matrix to dispersal (Balkenhol & Waits, 2009; Trombulak & Frissell, 2000). As populations become smaller and isolated, they might exhibit higher rates of inbreeding through genetic drift, resulting in an increased risk of population extinction (McCauley, 1991). Understanding the influence of LTIs on wildlife dispersal patterns is thus of critical importance to fuel conservation policies.

The most obvious detrimental effects of LTIs on dispersal success are direct collisions with vehicles and physical crossing impediment when infrastructures are for instance fenced (Forman & Alexander, 1998; Trombulak & Frissell, 2000). Most animals are affected, from small invertebrates to large mammals (Balkenhol & Waits, 2009; Fahrig & Rytwinski, 2009). LTIs may also induce behavioral alterations that further affect nearby populations (Trombulak & Frissell, 2000). For example, both breeding migrations and reproductive success of anurans can be perturbed by main roads due to sound interference with males mating calls (Bee & Swanson, 2007), in turn possibly impacting effective dispersal and thus gene flow (Ronce, 2007).

Over the past fifteen years, “molecular road ecology” has emerged as a fully-fledged discipline to thoroughly estimate landscape functional connectivity (Balkenhol & Waits, 2009; Holderegger & Di Giulio, 2010). Building on population genetics, landscape ecology and spatial statistic tools (Manel & Holderegger, 2013), its objective is to elucidate how the genetic variability is influenced by LTIs and other anthropogenic obstacles, with numerous applications in species management and conservation (Segelbacher et al., 2010). However, one major limitation of such studies is that they generally focus on a single species (Balkenhol & Waits, 2009; D. Keller et al., 2015), while different species may actually respond differently to the same type of infrastructure. Furthermore, they also often focus on a single LTI, while multiple LTIs are commonly built next to each other because of technical and economic constraints, notably within valleys or along coastlines: although the impacts of LTIs are then expected to add up and result in a “cumulative” barrier effect, some LTIs might actually be neutral to movement or even create corridors to dispersal (Bartzke et al., 2015), these antagonistic effects making the whole picture even more complex. For example, Paquet and Callagan (1996) showed that a motorway strongly impeded crossing events in wolves but that a railway and power lines located within the same study area together redirected wolves movements and thus rather acted as corridors. In the same vein, Latch et al. (2011) found that gene flow in the desert tortoise *Gopherus agassizii* was affected by roads but not by power lines. In highly fragmented landscapes, it is thus highly advisable to assess the concomitant influence of all existing LTIs using a multi-species approach to adopt efficient conservation policies (D. Keller et al., 2015; Richardson et al., 2016).

In this study, we assessed the respective and cumulative impacts of six French LTIs in four terrestrial species with contrasted life history traits (two vertebrates and two insects including a flying species) using molecular data. We hypothesized that flying species would be less affected by LTIs than ground-dwelling ones and that large infrastructures carrying vehicles (roads, motorways, railways) would overall be more impactful than infrastructures carrying energy (power lines, gas pipelines). We also hypothesized that the impacts of some LTIs might accumulate to shape spatial patterns of gene flow in studied species.

## 3. MATERIAL AND METHODS

### 3.1. Study area and biological models

The study was carried out in the ‘Périgord’ region in southwestern France (45°07’31.8’’N; 0°58’56.9’’E; Fig. 1). It is a 300 km^2^ rural landscape composed of limestone plateaus including crops, mowed meadows, deciduous forests and small villages. The hydrology is limited to small streams and ponds. Altitude ranges from 91 to 294 m above sea level. Six types of LTIs are present in this study area (from the widest to the narrowest): a fenced motorway (A89) commissioned in 2004; a low traffic single-track railway built in the 19^th^ century; a high traffic country road (D6089) present since the 18^th^ century; a power line and a gas pipeline constructed in 1962 and 1955, respectively, both associated with breaches in forest cover; a 1370 km dense network of low traffic secondary roads (Fig. 1).

**Figure 1:**
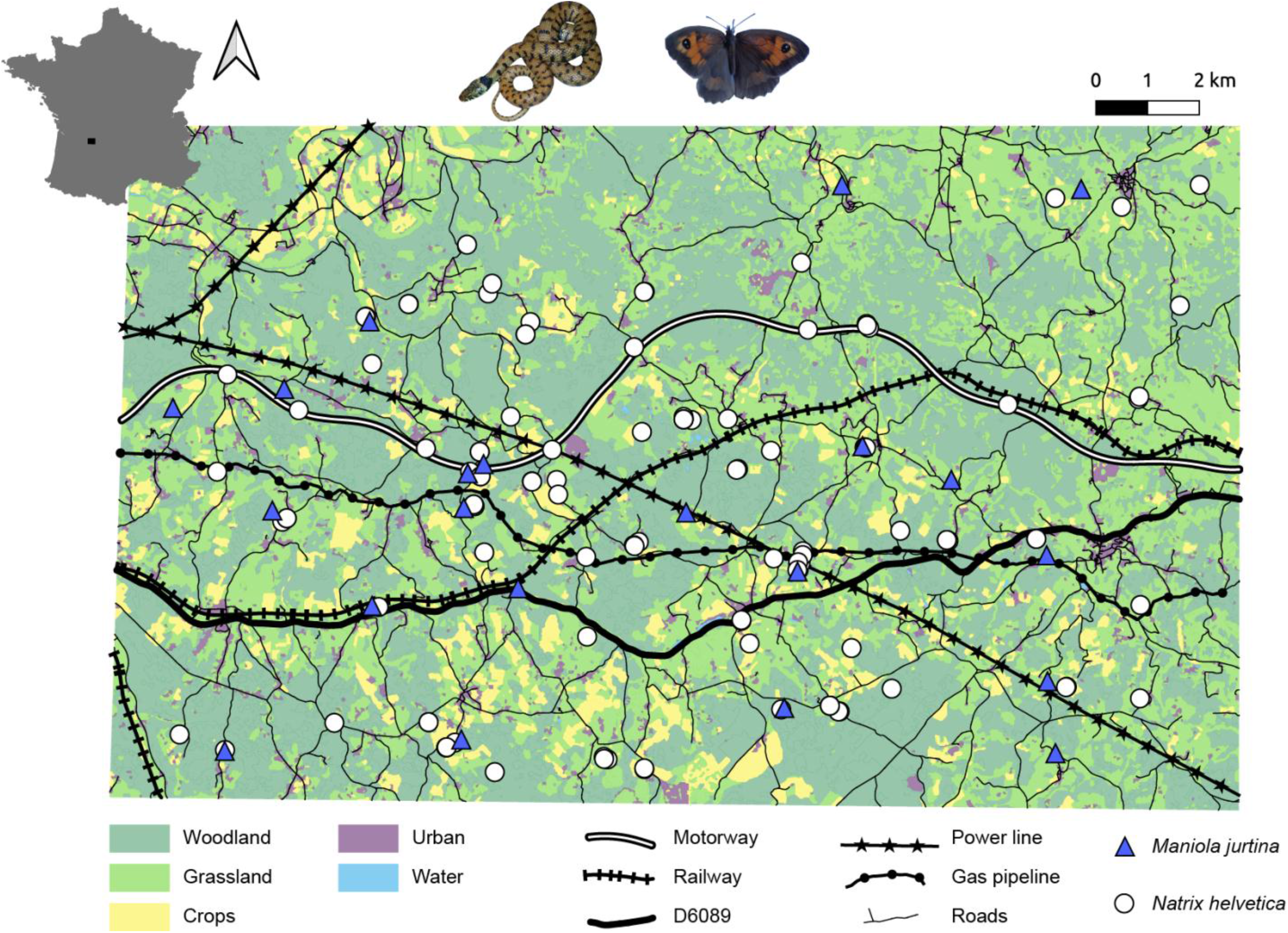
Study area in southwestern France and sampling locations of *Natrix helvetica* (115 individuals) and *Maniola jurtina* (21 populations of about 30 individuals each). For these two species, no genetic structure was identified (see result section).

We considered four species with various life history traits in order to span a large amount of biological variability: two vertebrates (the snake *Natrix helvetica* and the midwife toad *Alytes obstetricans*) and two insects (the butterfly *Maniola jurtina* and the ground-beetle *Abax parallelepipedus*). *Alytes obstetricans* is a small toad widely distributed in Western Europe. It is highly sensitive to fragmentation because local populations are known to function as relatively independent entities with strong genetic structuring (Tobler et al., 2013). *Natrix helvetica* is also widely distributed in Western Europe and is considered to exhibit good dispersal abilities, with individuals travelling over more than 1 km in less than a month (Pettersson, 2014). A previous study did not detect any genetic structure in this species in an intensively used agricultural landscape, indeed suggesting good dispersal ability in fragmented environments (Meister et al., 2010). *Maniola jurtina* is a univoltine butterfly which is very common in Europe with locally very high densities. It shows medium dispersal capacity with mean dispersal distances ranging from 50 to 300 m (Stevens et al., 2013). Previous studies revealed that both land cover (arable lands and forests) and LTIs (motorway and railway) could affect its dispersal (Remon et al., 2018; Villemey et al., 2016). Finally, *Abax parallelepipedus* is an opportunistic carnivorous ground-beetle that inhabits the upper layer of forest litter (Loreau, 1987). It typically shows limited dispersal capacity, avoids open habitats and is highly sensitive to fragmentation by roads (I. Keller et al., 2004).

### 3.2. Sampling and genotyping

All captures were authorized by Préfecture d’Aquitaine (ref number: AD_AD_150224_arrete_06-2015_terroiko). For all species, tissues were collected between April and September in 2015 and 2016. For the two vertebrate species *N. helvetica* and *A. obstetricans*, we followed an individual-based sampling design due to low abundances in the field. Individual-based sampling design has been proved to be a good alternative method to population-based sampling design as less individuals are required per sampling location (1 to 4) and more geographical locations can be sampled over the landscape (Luximon et al., 2014; Prunier et al., 2013). Accordingly, the entire study area was prospected to collect toads and snakes, at night and at day time, respectively. We mainly focused on sampling locations with high probability of presence such as wetlands, ponds, rivers, woodland edges and small villages. To attract snakes and facilitate data collection, 108 artificial shelters were also laid across the study area. When an individual was detected, it was hand-captured and manipulated directly in the field. A GPS location (Garmin Etrex20, USA) was recorded for each captured individual (see Fig. 1 and 2 for sampling locations). Each individual was sexed, measured, weighed, marked (to avoid sampling individual twice) and a genetic sample was collected. Captured toads were marked using 7×1.35 mm FDX-B Passive Integrated Transponder (PIT) tags (Loligo Systems, Denmark) and a non-destructive genetic sample was collected by gently opening mouth with a little metal spoon and swabbing mouth cavity for about 10 seconds (Broquet et al., 2007). We used ventral scales clipping following Brown and Parker (1976) to both mark snakes and collect DNA. We also opportunistically collected genetic samples from snakes and amphibians found dead (road kill or predation) and from snake shed skins.

**Figure 2:**
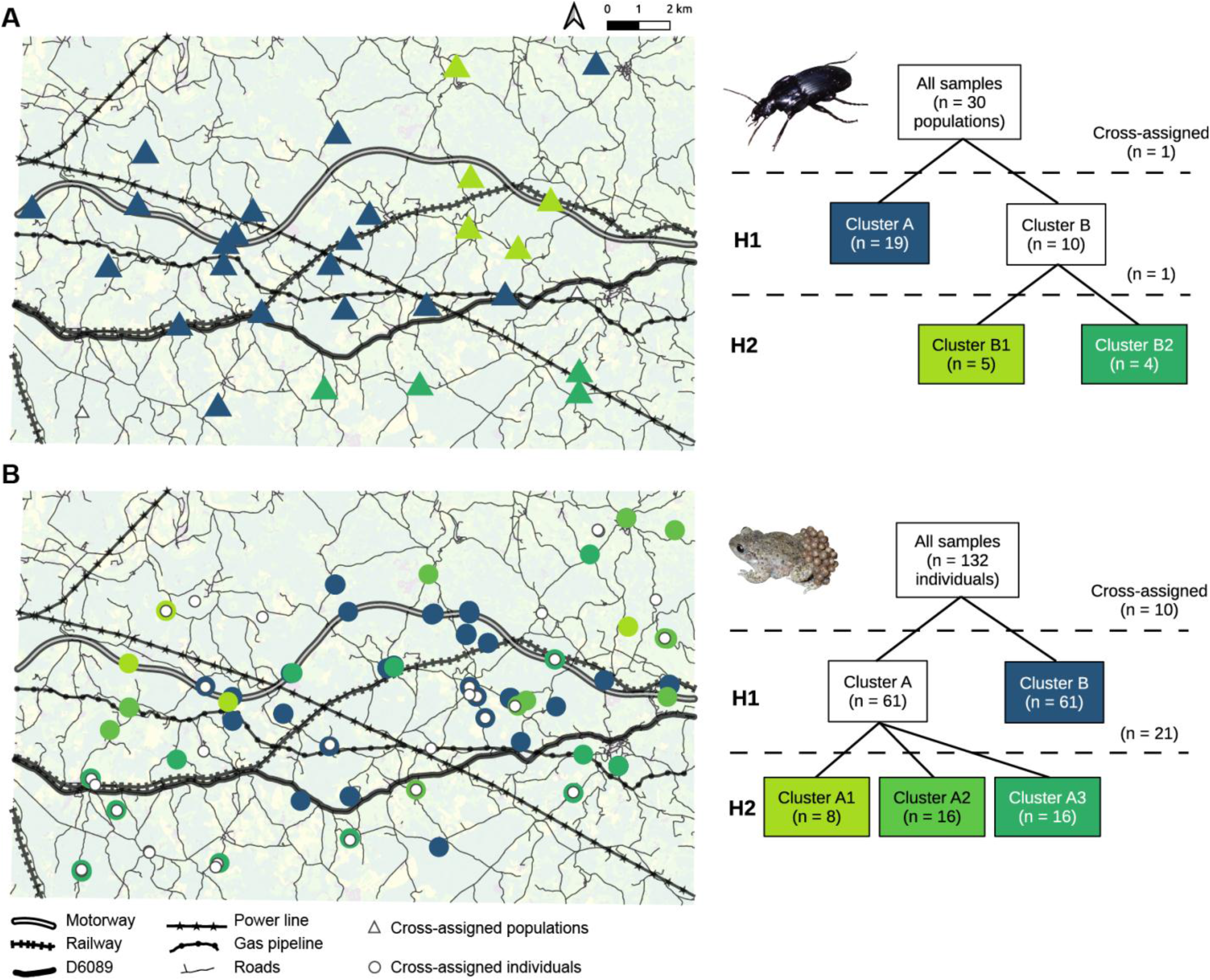
Left panels: STRUCTURE outputs for *A. obstetricans* (132 individuals in 56 sampling locations) and *A. parallelepipedus* (30 populations of about 30 individuals each) plotted over the study area. Right panels: hierarchical splits of inferred clusters from the first to the second hierarchical level. Each box represents a cluster, with n the corresponding number of assigned samples. The number of cross-assigned samples at each hierarchical level (Q-values < 0.6) is also indicated.

The two insect species *M. jurtina* and A. *parallelepipedus* were sampled within 30 sites using a classical population-based sampling design. Site locations were obtained by dividing the study area into 30 sectors using a 5×6 regular grid in QGIS (V. 2.8). In each sector and each species, a single sampling site was chosen according to the presence of suitable habitats (woodlands for beetles and grasslands for butterflies). At each sampling location, 30 individuals were sampled, resulting in 900 genetic samples per species (see Fig. 1 and 2 for sampling locations). Butterflies were captured during day time with nets. Beetles were trapped using non-lethal dry pitfalls. Pitfalls were 20 cm in diameter and 15 cm in depth and were arranged in circles at regular intervals of 5 m. They were emptied every day until 30 individuals were captured. For both insect species, we collected the middle right leg of each captured individual, as both a source of DNA and a way to avoid sampling the same individual twice.

All genetic samples were stored in 70 % EtoH until DNA extraction. All material for marking animals and collecting genetic samples was washed and disinfected using absolute ethanol between each individual sampling. Care was taken to minimise animal handling and stress and all individuals were rapidly released at the place of capture after manipulation. We amplified 13, 14, 15 and 14 polymorphic microsatellite loci in *N. helvetica, A. obstetricans, M. jurtina* and *A. parallelepipedus*, respectively. For a detailed procedure of DNA extraction, amplification and genotyping, see Appendix A. Some individuals could not be correctly genotyped because of insufficient amounts of DNA: genotypes with more than 2 loci presenting missing values were discarded to allow robust subsequent genetic analyses. We used Genepop 4.2 (Rousset, 2008) to test for linkage disequilibrium among pairs of loci and deviation from Hardy-Weinberg Equilibrium after sequential Bonferroni correction to account for multiple related tests (Rice, 1989). The presence of null alleles was tested using MICROCHECKER 2.2.3 (Van Oosterhout et al., 2004). Loci with null alleles and/or in linkage disequilibrium were discarded, resulting in the final selection of 13, 10, 6 and 10 microsatellite loci in toads, snakes, butterflies and beetles, respectively (Appendix A).

### 3.3. Genetic structure and genetic distances

The presence of related individuals in data sets may lead to an over-estimate of the number of clusters when assessing population structure and thus bias subsequent genetic analyses (E. C. Anderson & Dunham, 2008). We therefore used COLONY2 (Jones & Wang, 2010) to identify and discard siblings within our individual-based data sets (*N. helvetica* and *A. obstetricans*, Appendix B). In addition, because sites were unevenly sampled for toads, we only retained a maximum of three randomly picked genotypes per sampling location, following Prunier et al. (2013). In the population data sets, we only retained populations for which at least 15 genotypes were available. The final data sets comprised 848 genotypes (30 populations) in *A. parallelepipedus*, 508 genotypes (21 populations) in *M. jurtina*, 115 genotypes in *N. helvetica* (68 sampling locations) and 132 genotypes in *A. obstetricans* (56 sampling locations).

For each of the four final data sets, genetic structure was investigated using STRUCTURE 2.3.4 (Pritchard et al., 2000) with the admixture and the correlated allele frequency models and prior sampling location information when structure in the data was too weak. We followed a hierarchical genetic clustering procedure (Coulon et al., 2008). At each hierarchical level, we tested the number K of clusters from 1 to 10 and repeated analyses for each value 5 times. Runs were performed with a burn-in period of 50 000 and 50 000 subsequent Markov chain Monte Carlo (MCMC) repetitions. We also checked that the alpha value had stabilised before the end of the burn-in period to ensure algorithm convergence. If convergence was not reached, we used a burn-in period of 100 000 and 100 000 MCMC repetitions. We then used STRUCTURE HARVESTER (Earl & vonHoldt, 2012) to obtain deltaK statistics to infer the optimal K-value. We used this optimal K-value to perform 20 runs with a burn-in period of 200 000 and 200 000 MCMC repetitions. We finally compiled the ten best runs using CLUMPP (Jakobsson & Rosenberg, 2007) to obtain individual or population ancestry values (i.e., Q-values), measuring the level of admixture among the inferred genetic clusters. Each individual or population was assigned to the cluster for which the Q-value was higher than 0.6, following Balkenhol et al. (2014). We then repeated the analysis for each inferred cluster separately until no more structure was found in the data. For each hierarchical level, we used individual- or population-based Q-values to compute pairwise matrices of ancestry-based hierarchical genetic distances (HGD; Balkenhol et al., 2014). HGD were only calculated for species displaying a significant genetic structure. When more than one hierarchical level was detected, each hierarchical level (HGD1, HGD2) was considered separately. We also computed classical genetic distances (GD), using the Bray-Curtis (bc) percentage dissimilarity index for the individual-based data sets and Fst for the population-based data sets. While these classical genetic distances are well suited to detect surface elements affecting gene flow at a regional scale, HGD have been shown to allow a better detection of sharp genetic variations caused by linear elements such as LTIs (Prunier et al., 2017).

### 3.4. Multiple linear regressions and commonality analyses

Both classical and hierarchical genetic distances were tested against the six types of LTIs present in our study area, along with a number of covariates likely to affect patterns of genetic differentiation (isolation-by-distance IBD, difference in altitude and the following landcover features: water, crops, woodlands, grasslands and urban areas), although assessing the respective influence of these non-LTI features was not the main scope of this study (details about the effects of non-LTIs are provided in Appendix H). All LTIs but the secondary road network were coded into binary pairwise matrices, with 0 indicated that individuals/populations of each pair were on the same side and 1 indicated that they were on either side of the LTI. Because of the density of the secondary road network in the study area, this LTI (hereafter simply called Roads) was treated as other landcover features. Landcover features were defined by digitalizing the entire study area in QGIS (V. 2.8) using national maps and aerial photographs (National Geographic Institute, France). Every element of the landscape was classified into 49 habitat types of the EUNIS Habitat Classification System (Davies & Moss, 1999). Botanic field surveys were also performed in 2015 to confirm the affiliation of certain habitat types. We combined these 49 elements into six main landcover predictors (Appendix C): Water (stagnant water bodies, streams and rivers), Crops (intensive and non-intensive cultures), Woodlands (all types of forests), Grasslands (uncultivated open lands), Urban (villages, industrial sites, *etc*.) and Roads (all roads excluding small trails, motorway and D6089 country road). These six classes were each rasterised at a 1 m resolution using ARCGIS 10.2.2 and its SPATIAL ANALYST extension. Each raster was then used to create a resistance surface based on the spatial density of the corresponding element in the landscape. This procedure hypothesizes that a pixel covered with 100% (respectively 0%) of an unfavourable landscape feature would be 100% resistant (respectively permeable) to gene flow and thus avoids assigning arbitrary resistance values to landscape features. To do so, we overlaid a 20 m grid on each spatial class and calculated the percentage of the element in each grid. For each resistance surface, we rescaled pixel resistance values to range from 1 (null or extremely low densities) to 100 (the element covers the entire pixel) and the final rescaled resistance surface was used in CIRCUITSCAPE 4.0 (McRae et al., 2016) to compute pairwise effective distances between individuals or populations. The IBD pairwise matrix was similarly obtained by running CIRCUITSCAPE on a uniform resistance surface only composed of pixels of value 1. Finally, altitude pairwise matrices were computed as the absolute values of pairwise differences in altitude between sampling locations.

The local influence of landscape features may go unnoticed if all pairs of genetic distances are retained, as isolation-by-distance might take over the influence of landscape features, with strong implications in terms of biological interpretation of results (C. D. Anderson et al., 2010; D. Keller et al., 2013). We thus considered subsets of pairwise data by defining a maximum Euclidean distance threshold between sampling locations. Following Cayuela et al. (2019), this distance threshold was selected for each species and each metric of genetic distances (GD or HGD) as the neighbouring distance maximizing the model fit of a classical multiple linear model including all predictors (see Appendix D for details). For each species, we then explored the relationship between subsets of each type of genetic distances (GD or HGD) and the corresponding predictors using standard multiple linear regressions. The contributions of predictors to the dependent variables were assessed using commonality analyses (CA).

Commonality analysis is a variance partitioning procedure allowing the detection and the withdrawal of statistical suppressors that are responsible for a distortion of model estimates (beta weights β and confidence intervals), thus providing decisive support when trying to assess the reliability of model parameters in face of multicollinearity. It also allows isolating the unique contribution U of each predictor to the variance in the dependent variable (for more details about CA, see Appendix E and Prunier et al., 2015, 2017; Ray-Mukherjee et al., 2014). We performed model simplification by discarding predictors identified as statistical suppressors in an iterative way following Prunier et al. (2017; see Appendix F for details).

In each final simplified model, we assessed levels of collinearity among predictors using Variance Inflation Factors VIF (Dormann et al., 2013). Because pairwise data are not independent, the p-values inferred from simplified models could not be interpreted: we thus computed 95 % confidence intervals around regression estimates using a jackknife procedure, with 1000 replicates based on a random removal of 10 % of individuals/populations without replacement (Peterman et al., 2014). These confidence intervals were used to assess the significance of the predictors’ contributions to the variance in the corresponding genetic distances. We considered that a predictor was a robust contributor to the variance in the response variable as soon as the confidence interval about the corresponding β value did not include 0. A predictor with a positive β value was associated with an increase in the genetic distances and was interpreted as impeding gene flow. On the contrary, a predictor with a negative β was associated to a reduction in genetic distances and was thus interpreted as promoting gene flow (Jacquot et al., 2017). In the case of LTIs, this interpretation translates into two categories of LTI effects: LTI+ (impeding gene flow) and LTI-(promoting gene flow). In order to make unique contributions comparable across models, we finally computed the relative unique contribution (i.e., U/R^2^) of each predictor to the explained genetic variance in each model.

In order to summarise our main findings across species, we summed the relative contributions of LTIs *versus* non-LTIs predictors as well as the relative contributions of LTI+ *versus* LTI-effects within each model. We then averaged these contributions across models for each species and across species. To summarise our main findings across LTIs, we also averaged the relative unique contributions of LTI+ *versus* LTI-effects for each LTI and across LTIs. Non-significant predictors or predictors that were absent from final simplified models (including LTI-when only LTI+ was present, and vice versa) were given a relative contribution of 0. Results were plotted in the form of 100% stacked barplots.

## 4. RESULTS

### 4.1. Genetic structures

Structure outputs indicated a single genetic cluster in both *N. helvetica* and *M. jurtina*, suggesting high gene flow across the study area in these species. On the contrary, we found strong hierarchical genetic clustering in *A. obstetricans* and in *A. parallelepipedus* (Fig. 2). In toads, we identified two hierarchical genetic clusters. At the first level, one cluster (B) was surrounded by a second cluster (A) with no clear geographical boundaries explaining this pattern (Fig. 2). Ten individuals could not be assigned to any of these two clusters (cross-assigned individuals), suggesting some exchanges between these two clusters. At the second hierarchical level, only cluster A was further divided into three sub-clusters: A1, A2 and A3. These three sub-clusters were not separated by clear geographical patterns and a high number of individuals (21) were cross-assigned, again suggesting frequent exchanges among them.

We similarly identified two hierarchical clustering levels in beetles (Fig. 2). At the first level, 19 populations were assigned to cluster A and ten were assigned to cluster B. Cluster A included populations sampled mostly in the western part of the study area and north of the road D6089 (Fig. 2). One population at the extreme south-west could not be assigned to any of these two clusters (cross-assigned). Cluster B, was further divided into two sub-clusters at the second hierarchical level. Clusters B1 and B2 were separated by the D6089 and the gas pipeline, with B1 in the north comprising five populations and B2 in the south comprising four populations. At the second hierarchical level, only one population could not be assigned to any of these two clusters (cross-assigned). This population was located between the road D6089 and the gas pipeline, exactly in-between clusters B1 and B2.

### 4.2. Multiple linear regression and commonality analyses

The maximum Euclidean distances between sampling locations that optimized the amount of variance in classical and hierarchical genetic distances (variance explained by full regression models) ranged from 2800 to 3500m in individual-based data sets and from 4500 to 18500m in population-based data sets (Table 1; Appendix D). After simplification (Appendix F) and whatever the model, VIF ranged from 1.00 to 1.70 (Appendix G), suggesting little collinearity among retained variables (Dormann et al., 2013).

**Table 1:** Outputs of multiple linear regressions and additional parameters from commonality analyses (CA) for each species and for each type of data set. DV represents the dependent variable: classical genetic distances (GD; calculated either with the Bray-Curtis dissimilarity index (bc) or with Fst) and hierarchical genetic distances (HGD1 and HGD2 for first and second level of hierarchy, respectively). For each model, the model fit (R^2^) was estimated from reduced scale analyses, with a maximum distance threshold between pairs of individuals or populations (Dist.) ranging from 2800 to 18500m. In each model and for each retained predictor, we estimated the structure coefficient (rs), the beta weight (β), as well as unique (U), common (C) and total (T) contributions. The relative contribution of each predictor (% of R^2^) was computed as U/R^2^. Significance of a predictor’s contribution to the dependent variable was estimated using confidence intervals (CI-inf and CI-sup). A CI that included 0 was considered as a non-informative predictor (indicated in italic). The putative effect of significant LTI predictors on gene flow (Barrier or Corridor) is also provided.

| Species | DV | Dist. (m) | R<sup>2</sup> | Predictor | rs | $\beta$ | CI-inf | CI-sup | U | C | T | % of R<sup>2</sup> | Effect |
| --- | --- | --- | --- | --- | --- | --- | --- | --- | --- | --- | --- | --- | --- |
| *A. obstetricans* | GD(bc) | 3000 | 11.8% | IBD | 0.283 | 0.126 | 0.066 | 0.198 | 0.009 | 0.071 | 0.080 | 7.9 |  |
|  |  |  |  | Altitude | 0.212 | 0.098 | 0.052 | 0.140 | 0.008 | 0.037 | 0.045 | 6.5 |  |
|  |  |  |  | Woodlands | 0.190 | 0.145 | 0.091 | 0.191 | 0.018 | 0.018 | 0.036 | 15.3 |  |
|  |  |  |  | Roads | 0.214 | 0.113 | 0.062 | 0.153 | 0.009 | 0.037 | 0.046 | 7.9 | Barrier |
|  |  |  |  | D6089 | 0.110 | 0.091 | 0.043 | 0.142 | 0.008 | 0.004 | 0.012 | 6.6 | Barrier |
|  | HGD1 | 2400 | 10.8% | Woodlands | 0.151 | 0.100 | 0.037 | 0.172 | 0.010 | 0.013 | 0.023 | 8.8 |  |
|  |  |  |  | Crops | 0.225 | 0.185 | 0.099 | 0.254 | 0.032 | 0.018 | 0.051 | 30.1 |  |
|  |  |  |  | Roads | 0.221 | 0.159 | 0.100 | 0.203 | 0.024 | 0.025 | 0.049 | 21.8 | Barrier |
|  |  |  |  | Railway | 0.145 | 0.108 | 0.048 | 0.178 | 0.011 | 0.010 | 0.021 | 10.3 | Barrier |
|  | HGD2 | 3500 | 19.9% | Woodlands | 0.240 | 0.188 | 0.134 | 0.240 | 0.031 | 0.026 | 0.058 | 15.7 |  |
|  |  |  |  | Urban | -0.207 | -0.241 | -0.276 | -0.203 | 0.047 | -0.004 | 0.043 | 23.3 |  |
|  |  |  |  | Roads | 0.196 | 0.184 | 0.134 | 0.238 | 0.033 | 0.006 | 0.039 | 16.3 | Barrier |
|  |  |  |  | D6089 | 0.196 | 0.196 | 0.145 | 0.250 | 0.037 | 0.001 | 0.039 | 18.7 | Barrier |
|  |  |  |  | Motorway | -0.124 | -0.120 | -0.159 | -0.076 | 0.014 | 0.002 | 0.016 | 7.0 | Corridor |
| *N. helvetica* | GD(bc) | 2800 | 4.2% | Roads | -0.109 | -0.125 | -0.193 | -0.062 | 0.015 | -0.003 | 0.012 | 35.4 | Corridor |
|  |  |  |  | Motorway | 0.125 | 0.148 | 0.078 | 0.221 | 0.021 | -0.005 | 0.016 | 50.8 | Barrier |
|  |  |  |  | Railway | -0.106 | -0.088 | -0.155 | -0.022 | 0.008 | 0.004 | 0.011 | 18.6 | Corridor |
| *M. jurtina* | GD(Fst) | 5500 | 19.9% | IBD | 0.209 | 0.264 | 0.001 | 0.490 | 0.066 | -0.023 | 0.044 | 33.3 |  |
|  |  |  |  | Woodlands | 0.306 | 0.315 | 0.077 | 0.519 | 0.089 | 0.004 | 0.093 | 44.8 |  |
|  |  |  |  | Power line | -0.266 | -0.180 | -0.388 | 0.046 | 0.030 | 0.040 | 0.071 | 15.3 |  |
| *A. parallelepipedus* | GD(Fst) | 6500 | 25.9% | Altitude | 0.103 | 0.121 | -0.023 | 0.251 | 0.015 | -0.004 | 0.011 | 5.6 |  |
|  |  |  |  | Grasslands | 0.494 | 0.498 | 0.372 | 0.610 | 0.248 | -0.004 | 0.244 | 95.9 |  |
|  | HGD1 | 18500 | 17.2% | Roads | 0.337 | 0.262 | 0.170 | 0.350 | 0.063 | 0.051 | 0.114 | 36.5 | Barrier |
|  |  |  |  | D6089 | 0.331 | 0.254 | 0.159 | 0.338 | 0.059 | 0.051 | 0.110 | 34.1 | Barrier |
|  | HGD2 | 4500 | 26.8% | Altitude | 0.230 | 0.223 | 0.056 | 0.397 | 0.049 | 0.004 | 0.053 | 18.3 |  |
|  |  |  |  | D6089 | 0.393 | 0.350 | 0.184 | 0.500 | 0.114 | 0.040 | 0.154 | 42.7 | Barrier |
|  |  |  |  | Motorway | -0.163 | -0.114 | -0.273 | 0.041 | 0.012 | 0.015 | 0.027 | 4.5 |  |
|  |  |  |  | Gas pipeline | 0.268 | 0.225 | 0.070 | 0.368 | 0.049 | 0.022 | 0.071 | 18.4 | Barrier |

When considering classical genetic distances in toads, the multiple linear regression explained 11.8% of variance (Table 1). Both the D6089 (U = 0.008) and Roads (U = 0.009) were associated with an increase in genetic distances in this model, thus suggesting barrier effects. They together uniquely contributed to 14.4% of explained variance. When considering the first level of hierarchical genetic distance (HGD1), the model explained 10.8% of the variance. Both Roads (U = 0.024) and Railway (U = 0.011) were associated with an increase in genetic distances (positive β values), here again indicating barrier effects. At the second level of the hierarchy, the model explained 19.9% of variance in HGD2. Roads and D6089 were, again, associated with an increase in genetic distances (positive β values) but the Motorway was also detected has having a weak but significant positive effect on toads’ effective dispersal (U = 0.014). When relative contributions of LTIs were summed in each model and then averaged across models, LTIs accounted for 46.7% of total explained variance (Fig. 3A). Infrastructures were mostly associated with an increase in genetic distances, with 92.1% of variance explained by LTIs stemming from the barrier effects (Fig. 3B) of the D6089, the Railway and Roads. The 7.9% left were explained by a reduction in genetic distances across the Motorway at the second level of the hierarchy (HGD2; Fig. 3B).

**Figure 3:**
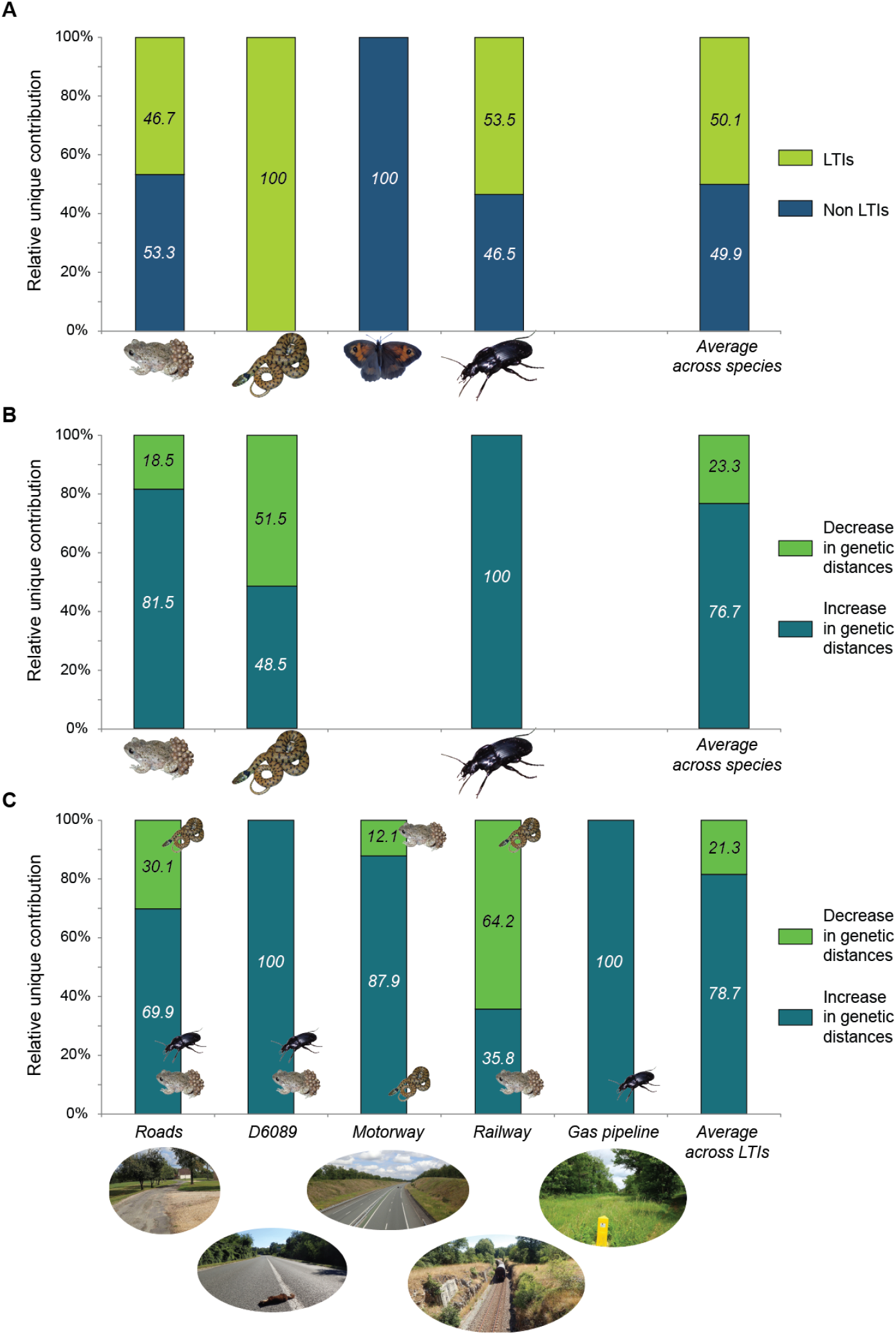
Stacked barplots of the averaged relative contributions of various predictors to the explained genetic variance. Panel A: For each species and for all species combined, averaged relative contributions of LTIs (all infrastructures combined) versus non-LTI predictors. Panel B: For each species (except *M. jurtina*) and for all species combined, averaged relative contributions of LTIs associated with an increase (barrier effect) versus a decrease (corridor effect) in genetic distances. Panel C: For each LTI (except the power line) and for all LTIs combined, averaged relative contributions of LTIs associated with an increase (barrier effect) versus a decrease (corridor effect) in genetic distances.

In snakes, the simplified model explained a small amount (4.2%) of variance in the dependent variable (Table 1) but only comprised LTIs predictors (Fig. 3A). The Motorway was associated with an increase in genetic distances and uniquely accounted for 48.5% of explained variance (U = 0.021; Fig. 3B). The two other LTIs (Roads and Railway) had unique contributions of 0.015 and 0.008, respectively, and both were associated with a reduction in genetic distances in this species, together accounting for 51.5% of explained variance (Fig. 3B).

In butterflies, the simplified model explained 19.9% of variance in Fst values (Table 1). The only LTI that remained in the final model was the Power line but it did not significantly contribute to the model predictive power. The entire genetic variability in this species was thus explained by IBD and Woodlands, both impeding gene flow (Fig. 3A and Appendix H).

In the ground-beetle, the simplified model explained 25.9% of the variance in Fst values (Table 1). The entire genetic variability was yet here explained by non-LTI features (Fig. 3A; Appendix H). When considering the first and the second level of the inferred hierarchical genetic structure, simplified models explained 17.2% and 26.8% of the variance in HGD1 and HGD2, respectively. In both cases, the D6089 was associated with an increase in genetic distances, indicating a consistent barrier effect (U = 0.059 in HGD1 and 0.114 in HGD2). In addition, Roads (HGD1) and the Gas pipeline (HGD2) were also detected as having negative effects on gene flow (U = 0.063 and 0.049, respectively). In HGD2, the Motorway did not significantly contribute to the model predictive power. Overall, explained variance in genetic distances was accounted for by both LTIs (53.5%) and non-LTIs elements (46.5%; Fig. 3A), with all LTIs being associated with an increase in genetic distances (Fig. 3B).

### 4.3. Assessment of infrastructure effects

Overall, 50.1% of the explained variance in genetic distances across all species was due to LTIs (Fig. 3A), of which 81.2% was associated with an increase in genetic distances, that is, with a barrier effect (Fig. 3B-C). The only LTI that did not contribute to genetic distances in any species was the Power line. On the contrary, the D6089 and the Gas pipeline were both systematically associated with barrier effects, in toads and beetles for the former and in beetles only for the latter. Other LTIs however showed more nuanced impacts, with corridor effects detected in some species (18.8% of explained variance by LTIs). While 87.9% of the overall genetic variability explained by the Motorway across species corresponded to a barrier effect in snakes, the remaining 12.1% corresponded to a reduction in genetic distances in toads (Fig. 3C). It was the opposite in the case of the railway, with 35.7% corresponding to a barrier effect in toads but 64.3% to a reduction in genetic distances in snakes. Finally, Roads acted as a barrier to gene flow in both toads and beetles (70.0%) but as a corridor in snakes (30.0%; Fig. 3C).

## 5. DISCUSSION

The goal of this study was to assess landscape functional connectivity in four species occupying a landscape fragmented by multiple LTIs. We were particularly interested in the potential cumulative or on the contrary the antagonistic effects of six LTIs. We used individual- and population-based regression analyses along with commonality analyses over restricted spatial scales to thoroughly evaluate the relative contribution of various landscape predictors to the variance in both classical and hierarchical genetic distances. We notably showed that LTIs were overall responsible for half of observed genetic variability across species but that the response of organisms to these LTIs was highly species-dependant. Most importantly, we found that LTIs did not only act as barriers to gene flow but might on the contrary promote gene flow, with some antagonistic effects across species.

Overall, LTIs were found to have a strong influence (either positive or negative) on gene flow, accounting for 50.1% of the total explained genetic variability across species and genetic distances. All ground-dwelling species were affected by LTIs, with contributions to the variance by LTIs ranging from about 50% in toads and beetles to 100% in snakes, contrary to the flying species *M. jurtina* whose genetic variability was only affected by distance and woodlands, as expected from a previous study (Villemey et al., 2016). Although butterflies have a lower probability to be impacted by vehicles than ground-dwelling species, previous studies showed that roads and motorways could hinder crossing events in this species (Polic et al., 2014; Remon et al., 2018). A direct Mark-Release-Recapture survey conducted in the same study area notably found that the motorway was responsible for a six-fold decrease in crossing events when compared to adjacent habitats (Remon et al., 2018). It is possible that large population sizes in *M. jurtina* are responsible for a temporal inertia in the setting-up of genetic differentiation since the creation of the motorway in 2004 (Landguth et al., 2010), but this study showed that some butterflies were able to cross it, thus possibly ensuring sufficient gene exchange across the landscape. Although we could not ascertain the negative aftermaths of human-induced fragmentation in *M. jurtina* from our genetic data, our study highlights the potential benefits of combining landscape genetics and Mark-Release-Recapture surveys (Cayuela et al., 2018).

As expected, LTIs were mainly associated with a reduction in gene flow, barrier effects accounting for 81.2% of the variance explained by LTIs across ground-dwelling species. LTIs carrying vehicles (roads, motorway and railway) were more impacting than infrastructures carrying energy (gas pipeline and power line). Roads and D6089 were responsible for most of inferred barrier effects in this landscape, with negative effects on gene flow in both toads and beetles. The motorway and the railway also accounted for non-negligible amounts of explained genetic variability but to a lesser extent than roads, only negatively affecting snakes and toads, respectively. In contrast, the contributions of LTIs carrying energy were less important. The gas pipeline negatively affecting gene flow in the ground-beetle only, probably in response to associated breaches in forest cover (Charrier, 1997), and the power line did not affect any studied species. These results suggest that conservation measures should primarily focus on infrastructures carrying vehicles rather than on infrastructures carrying energy (Bartzke et al., 2015), although we acknowledge that some taxa not considered in this study, for instance birds, might be negatively affected by LTIs such as power lines (Loss et al., 2015).

Despite these general negative impacts of LTIs on gene flow, we found that species showed very different responses to the same LTI, which perfectly highlights the importance of considering functional rather than just structural landscape connectivity in empirical studies (Taylor et al., 2006). Three of the six studied LTIs were associated with an increase in genetic distances in toads, these barrier effects together accounting for 92.1% of genetic variance explained by LTIs. Roads and D6089 were the main barriers to dispersal in *A. obstetricans*, affecting both classical (GD) and second-order hierarchical genetic distances (HGD2). In addition, Roads also impeded gene flow at the first level of the hierarchy (HGD1). Garcia-Gonzalez et al. (2012) similarly found that all roads, including small secondary roads, acted as barriers to gene flow in *A. obstetricans* in northern Spain. Amphibians are particularly vulnerable to road kills because of their numerous movements during dispersal but also during seasonal migrations between breeding water bodies and shelters (Fahrig & Rytwinski, 2009). Although these results advocate for effective mitigation measures to limit road kills of amphibians (Beebee, 2013), it is important to keep in mind that other road features such as traffic noise may also affect amphibians population dynamics (Bee & Swanson, 2007).

In addition to toads, we found that roads also deeply impacted the ground-beetle *A. parallelepipedus*, a result congruent with Keller et al. (2004). Roads and D6089 explained the whole genetic variance at the first hierarchical level (HGD1) resulting in clusters A and B (Fig. 2A). At the second hierarchical level (HGD2), the D6089 (but also the gas pipeline) was associated with the split of cluster B into two sub-clusters (Fig. 2A) and thus probably further impacted gene flow. Roads may act as barrier to gene flow because of road kills but also because ground-beetles may be reluctant to cross roads due to behavioural changes (Holderegger & Di Giulio, 2010).

Contrary to roads, we found that the motorway and the railway showed limited barrier effects. The only species that was negatively affected by the motorway was the snake *N. helvetica*. We here revealed that half of the explained genetic variability in snakes resulted from the negative impacts of the motorway. Because it is fenced with fine mesh, snakes can only reach the other side by using crossing structures (bridges, underpasses, culverts, etc.). These crossing structures may yet be seldom used by snakes due to inadequate placement, architectural design and snakes’ behaviour (Woltz et al., 2008). Thermoregulatory behaviour of reptiles is probably the main reason why individuals would not use underpasses, as a 50 m-length underpass would provide inadequate thermal conditions due to the absence of sunlight. In addition, Baxter-Gilbert et al. (2015) evaluated the effectiveness of different mitigation measures implemented to reduce reptile road mortality (including underneath culverts) and found that these structures were seldom used by reptiles. Underpasses may yet be used by other taxa such as amphibians and insects (Georgii et al., 2011), which may explain why the motorway was only found as acting as a barrier in a single species.

Similarly, only one species was negatively affected by the railway. At the first level of the hierarchy (HGD1), we found that a positive relationship between the presence of the railway and genetic distances in toads although clusters A and B were not clearly separated by this LTI, suggesting a modest barrier effect. Railways are known to restrict gene flow in some amphibian species such as frogs or salamanders and many studies on train collision with wildlife reported a high abundance of amphibian kills, representing up to 47% of all vertebrate records (Borda-de-Água et al., 2017). However, the railway in our study area had a low traffic density with approximately 10 trains/day, and train collisions may not be the only driver of the observed reduction in gene flow in *A. obstetricans*. The physical features of the railway are more likely to explain this pattern. Amphibians indeed have a high probability to be trapped between or along rail tracks, making them more vulnerable to both collisions and desiccation than other vertebrates (Budzik & Budzik, 2014). The studied railway was more than 150 years-old, which seems to be of sufficient duration for the detection of a barrier effect from genetic data (Landguth et al., 2010; Prunier et al., 2014) and suggests that this LTI was actually permeable to the movement of other species.

Our most striking finding is that, instead of acting as barriers, some LTIs might somehow promote dispersal. This “corridor effect” accounted for 18.8% of the overall genetic variance explained by LTIs across species and concerned both vertebrates. We first found that, at the second level of the hierarchy (that is, at a more local scale), gene flow in toads was promoted by the motorway. This counter-intuitive genetic pattern could stem from the availability of new habitats provided by the LTI. Adults and tadpoles of *A. obstetricans* were indeed detected in eight out of the ten storm-water retention ponds present along the studied motorway (data not shown). These ponds may provide favourable breeding habitats, free of predatory fish and surrounded by sand or gravel, the ideal substrates to build their burrows. Furthermore, the motorway is crossed by underneath culverts and tracks which are good dispersal corridors for amphibians (Georgii et al., 2011), especially when they are filled with water. This is not the first study showing a potential positive effect of a motorway on amphibian gene flow. Prunier et al. (2014) revealed that a 40-years old motorway was not a barrier for the alpine newt (*Ichthyosaura alpestris*) and could even serve as a longitudinal dispersal corridor when the surrounding landscape matrix is highly unfavourable. Interestingly, they even found negative relationships between genetic distances and presence of the motorway, indicating that, as in our study, gene flow across the motorway was probably enhanced; but because they analysed their data using one-tailed Mantel tests, they did not discuss this possibility. These results might yet be interpreted with caution due to the recent age of the motorway (<15 years old): this genetic pattern could stem from ancestral landscape configurations and direct monitoring surveys are now necessary to confirm that the motorway is indeed not an obstacle for toads.

Despite limited explained variance in snakes, we also identified two LTIs possibly acting as corridors in this species, Roads and Railway, together accounting for 51.5% of genetic variance explained by LTIs. Roads are known to be responsible for a high mortality in snakes (Rosen & Lowe, 1994): they bask on road surfaces to absorb radiant heat but this behaviour increases the probability of collisions and can result in a reduction in gene flow across roads (Clark et al., 2010). However, we found the exact reverse pattern, with Roads associated with a reduction in genetic distances in *N. helvetica*. This conflicting result can be explained by an attractive effect of roads and road verges that provide basking surfaces, reinforced by a limited traffic volume in our study area. In addition, the distribution of grass snakes being strongly dependent on wetlands for foraging, water-filled ditches often found alongside secondary roads may provide rich feeding areas, resulting in a local increase in snake abundance that favours road crossings and gene flow: a similar explanation was proposed by Johansson et al. (2005) who found a positive effect of gravel roads and associated ditches in the common frog (*Rana arvalis*). The railway was probably as attractive as Roads for snakes, which may similarly explain gene flow enhancement observed in snakes. Railway embankments provide important alternative habitats for reptiles with optimal thermal conditions for basking (Borda-de-Água et al., 2017). Even active lines can harbour particularly high diversity in reptile species, notably because human presence is scarce and because reptiles can perceive vibrations transmitted through the rail tracks and the ballast when a train approaches, allowing them to reach a shelter before collision (Borda-de-Água et al., 2017).

## 6. CONCLUSION

The accumulation of LTIs within landscapes is emerging as an important concern and local conservation policies are to be fuelled by a thorough assessment of landscape functional connectivity. Although focusing on a single species may help corridor planning (Baguette et al., 2013), we here illustrated how important it is to assess landscape connectivity from a multi-species perspective. Overall, we did not find consistent evidence of a cumulative barrier effect of the six LTIs across species: indeed, butterflies were not influenced by any LTI and snakes were only negatively impacted by the motorway. The case of toads and beetles was yet much more compelling. These two species were the most heavily impacted, with patterns of gene flow affected by various LTIs at different spatial scales. Roads were critical determinants of gene flow across all hierarchical levels in both species, but the railway and the gas pipeline respectively reinforced these impacts in *A. obstetricans* at the first hierarchical level and in *A. parallelepipedus* at the second one. In these two species, the impact of the accumulation of LTIs was thus more a question of a hierarchical than of a cumulative effect of barriers. Importantly, we also showed that some LTIs, acting as barrier for some species, could somehow promote gene flow in some others, leading to antagonistic LTIs effects: the motorway may have affected snakes but may have also provided favourable habitats for toads, while the railway may have affected toads but may have also provided favourable habitats for snakes. Considering the high variability in species response to LTIs, we argue that considering a single species may lead to counterproductive mitigation measures and that integrative approaches based on multiple species are to be more systematically considered. As it obviously seems impossible to assess functional connectivity in all existing species in a given landscape, it is also necessary to determine the extent to which species-specific mitigation measures can benefit the largest number of species, and, more generally, to investigate which life-history traits drive the taxonomic-specific response of organisms to the presence of LTIs.

## Acknowledgements

We gratefully thank M. Guillau, N. Macel, A. Dubois, E. Languille, T. Langer, D. Jacquet, A. Mira, E. Garcia, R. Roudier, A. Bideau, A. Brisaud, E. Chevallier, L. Tillion, M. Sanders, K. Henderson, A. Verzeni and O. Berggreen for their help in fieldwork. This study was granted by the French Ministry of Ecology, Energy, Sustainable Development and the Sea (CIL&B-ITTECOP-FRB Program).

## APPENDICES

A. Laboratory procedures and microsatellite markers

B. Sibship reconstruction

C. Landscape features defining the six main retained landscape

D. Spatial scale of analyses

E. Commonality analyses

F. Intermediate steps of commonality analyses on vectors

G. Correlations among final predictors

H. Supplementary results and discussion

I. References for appendices

### A. Laboratory procedures and microsatellite markers

For all species, we used a Qiagen Type-it Microsatellite kit. We extracted total DNA from insect legs, scales and swabs using the DNeasy Blood and Tissue kit (Qiagen, Valencia, CA). Before enzymatic digestion, each insect leg and scale was cut in 4-6 pieces to facilitate DNA extraction. Buccal swabs were used as is. For *Natrix helvetica* and *Alytes obstetricans*, we amplified 13 (Pokrant et al. 2016) and 14 (Tobler et al. 2013, Maia-Carvalho et al. 2014) polymorphic microsatellite loci, respectively. For both species, loci were amplified in 10 µl reaction volumes containing 2 µl multiplex PCR Master Mix, 1.2 to 1.6 µl of primer mix (between 0.13 and 0.25 µM of each primer), 5.4 to 5.8 µl of purified water and 1 µl of template DNA (10-20 ng µl-1). For *Maniola jurtina*, we amplified 15 polymorphic microsatellite loci (Richard et al. 2015) in three Multiplexes, in 10 µl reaction volumes containing 2 µl multiplex PCR Master Mix, 0.7 µl of primer mix (between 0.03 and 0.08 µM of each primer), 4.3 µl of purified water and 3 µl of template DNA (1-10 ng µl-1). For *Abax parallelepipedus*, we amplified 14 polymorphic microsatellite loci (Marcus et al. 2013) in three Multiplexes, in 5 µl reaction volumes containing 1 µl multiplex PCR Master Mix, 0.7 µl of primer mix (between 0.04 and 0.11 µM of each primer), 2.3 µl of purified water and 1 µl of template DNA (approx. 10 ng µl-1).

Polymerase Chain Reaction (PCR) conditions were set on an Applied Biosystems thermal cycler. For the two vertebrate species, conditions were set as follows: initial denaturation 10 min at 95°C; 30 cycles of 30 s at 95°C, 90 s at 51 to 60°C (depending on the multiplex) and 30 s at 72°C; final elongation of 5 min at 72°C. For the two insect species, conditions were set as follows: initial denaturation 10 min at 94°C; 40 cycles of 30 s at 94°C, 90 s (for the 10 first) or 30 s (for the 30 following) at 61°C (*A. parallelepipedus*) or 56°C (*M. jurtina*) and 30 s at 72°C; final elongation of 5 min at 72°C. All PCR products were ten times diluted and were run on an ABI 3730 DNA Analyser (Applied Biosystems) with the GeneScan-600 LIZ size standard. Genotyping was performed with GENEMAPPER 5.0 (Applied Biosystems) and all peaks were manually confirmed.

In the *A. obstetricans* data set, there was no evidence of linkage disequilibrium among loci. We found evidence of null alleles for locus Aly7. Accordingly, we retained 13 loci for subsequent analyses (Aly28, Aly3, Aly4, Aly17, Aly19, Aly20, Aly23, Aly24, Aly25, Aobst14, Aobst15, Aobst16 and Aobst17).

In the *N. helvetica* data set, two loci could not be amplified (Nsµ3 and 3TS) either in multiplex or in standalone PCR. There was no evidence of null alleles, but we found evidence of linkage disequilibrium between loci Natnat05 and µNt8new and between loci Natnat05 and TbuA09. Therefore, we only retained 10 loci for subsequent analysis (Natnat09, µNt8new, µNt3, µNt7, µt06, Natnat11, Eobµ1, Eobµ13, TbuA09 and 30).

In the *M. jurtina* data set, the locus Mj2410 was discarded as it showed sex linkage (Richard et al. 2015, Villemey et al. 2016). As Villemey et al. (2016), we found evidence of frequent null alleles for loci: Mj5522, Mj5287, Mj5647, Mj3956, Mj5563, Mj0272, Mj0283 and Mj3637. Thus, we only retained six loci for subsequent analysis (Mj0008, Mj7132, Mj0247, Mj7232, Mj4870 and Mj5331).

In the *A. parallelepipedus* data set, there was no evidence of linkage disequilibrium among loci. We found evidence of null alleles for loci: apar14, apar44, apar46 and apar50. Then, we retained 10 loci for subsequent analysis (apar20, apar50, apar27, apar34, apar32, apar12, apar23, apar25, apar02, apar46, apar05, apar44, apar14, apar06).

The following tables describe the specificity of the microsatellite markers tested for the four species followed in this study. Gray colours represent markers that were not used in the landscape genetic analyses either because they could not be amplified, showed sex-linkage, presence of null alleles or linkage disequilibrium.

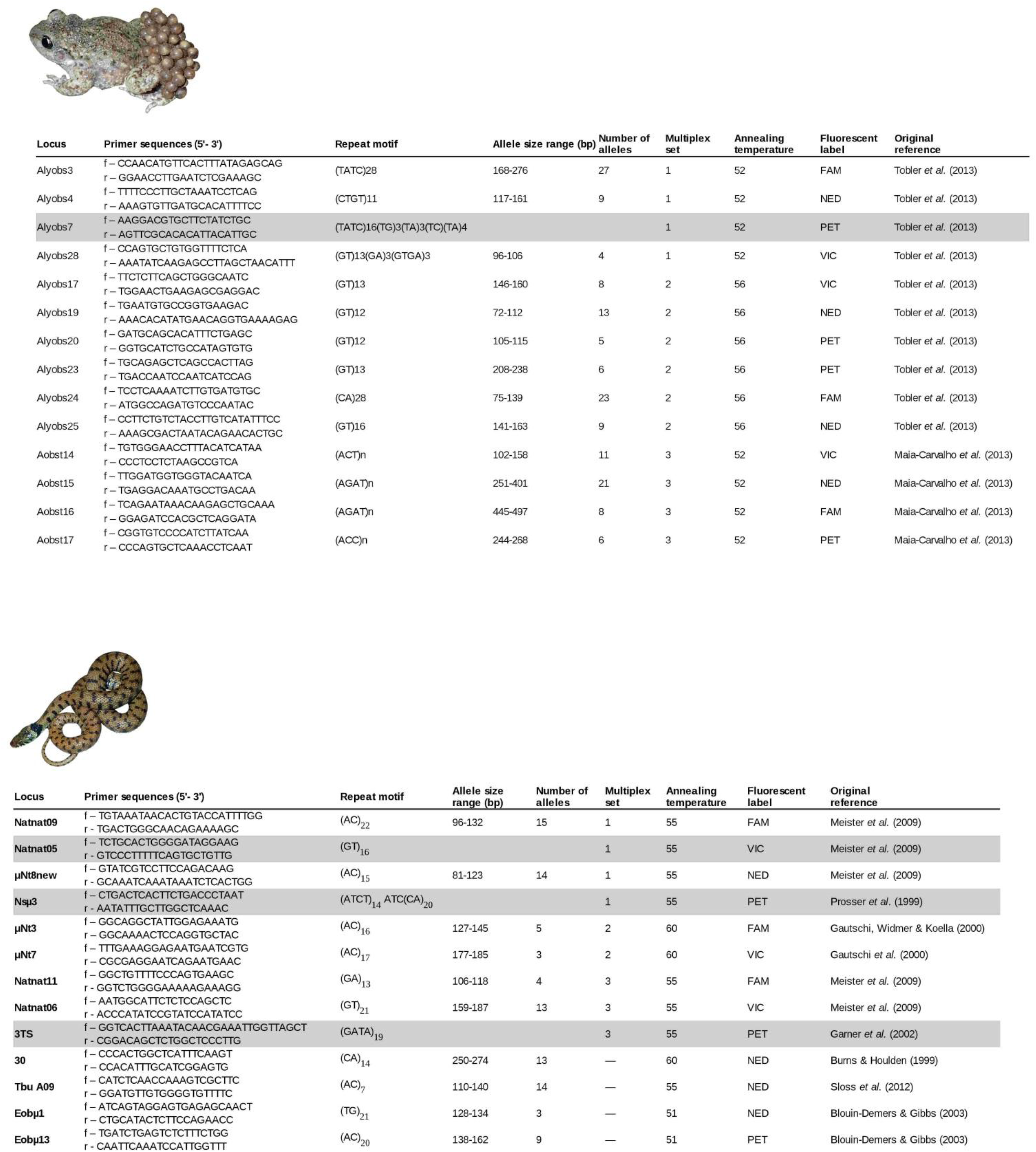
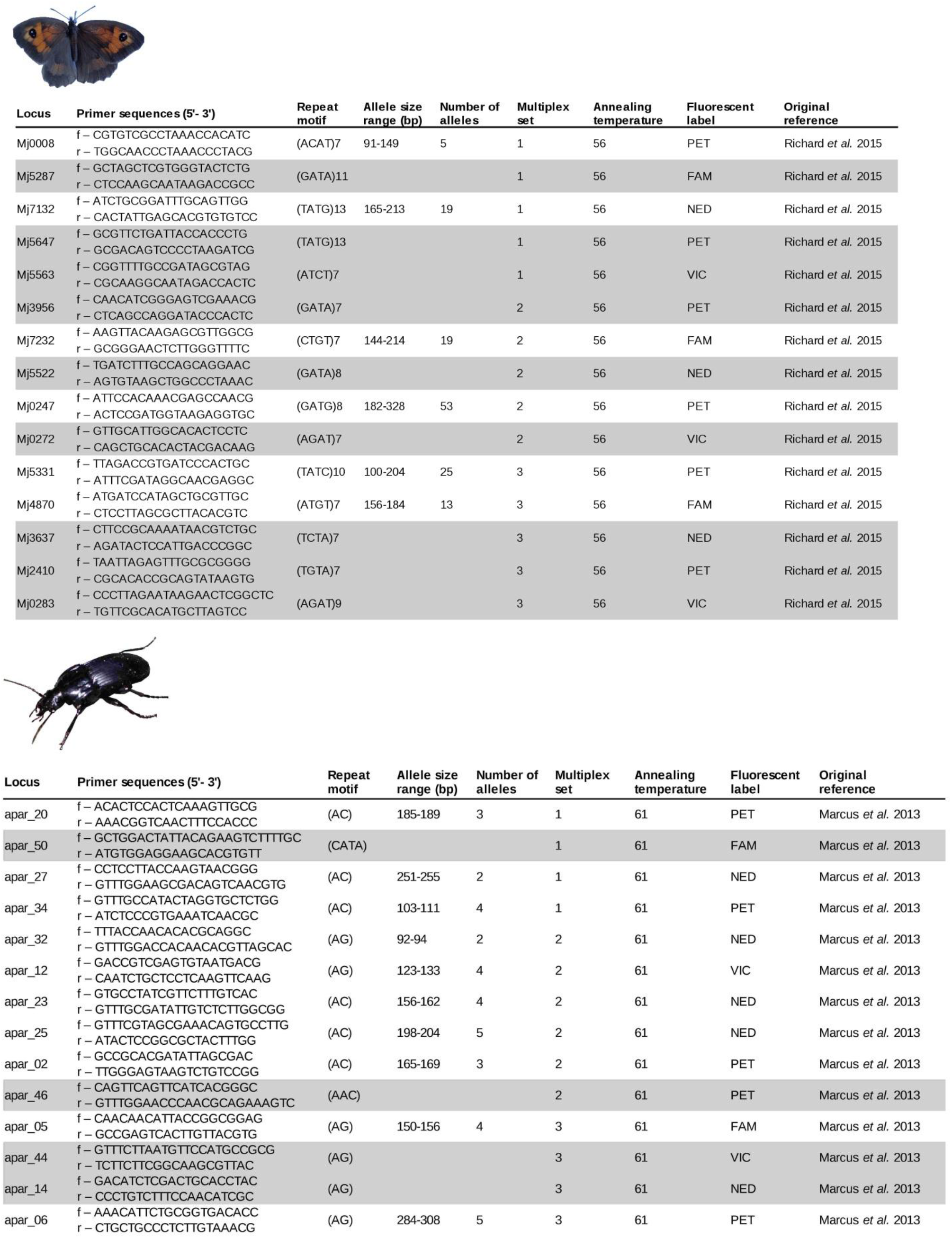

### B. Sibship reconstruction

We used COLONY2 (Jones and Wang 2010) to identify full-sib and parent-offspring groups among our individual-based data sets (*N. helvetica*) and (*A. obstetricans*). We used the full-likelihood approach based on the individual multilocus genotypes. For both species, we assumed that males and females were polygamous (for the snake, see Meister at al. 2012a). All individuals were considered as potential offspring and no a priori candidate parental genotype was defined. Allele frequencies were determined directly from genetic datasets. We ran three independent long runs with various seed numbers to test for congruence among results. Only relationships with an associated inclusion probability higher than 95% were considered as significant. In each group of related individuals, we randomly retained one genotype. Accordingly, 76 genotypes from *A. obstetricans* and one genotype from *N. helvetica* (corresponding to the shed skin of an already sampled individual) were discarded.

### C. Landscape features defining the six main retained landscape

- Water: Stagnant water; Streams; Ditches; Rivers
- Crops: Intensive monocultures; Gardens; Orchards; Vineyards; Vegetable gardens or horticultures
- Woodlands: Recent logged forests; Coniferous forests; Decideous forests; Riparian forests; Mixed woodlands; Heathlands; Hedgerows; Tree plantations; Bushlands
- Grasslands: Grass stripes; Forest clearings; Openings; Grazed pastures; Dry grasslands; Hayed meadows; Meadows; Trails and paths; Rocky lands; Abandoned arable lands
- Urban: Agricultural buildings; Residential Buildings; Waste disposals; Electric pylons; Water tanks; Artificial gardens; Domestic gardens; Cemeteries; Sport equipements such football fields; Surroundings of agricultural buildings; Campings; Car parks; Greenhouses; Open cast mines; Stone quarry; Industrial sites; Urban paved areas; Windmills
- Roads: Gravelled roads; Paved roads

### D. Spatial scale of analyses

The spatial scale retained in landscape genetic analyses can deeply influence the conclusions of studies (Keller et al. 2013, Schregel et al. 2018). The local influence of landscape elements on genetic distances can remain unnoticed if spatial scale retained is wide in comparison to dispersal capacities of individuals (Anderson et al. 2010). Accordingly, we did not use all possible pairs of populations or individuals in our data sets. For each dataset, we retained a subset of pairwise data by defining a maximum Euclidean distance between pairs, following Cayuela et al. (2019). The maximum Euclidean distance was selected as the neighbouring distance maximizing the R^2^ of our full model including all predictors in a classical multiple linear regression. This retained distance was higher than the minimum distance in a neighbouring graph which ensured that no individual was excluded from the network (Jombart et al. 2008). It was estimated using Gabriel graphs with the “adegenet” package (Jombart 2008) in R 3.3.2 (R Core Team, 2015). Subsequent analyses were only run with pairwise data associated with Euclidean distances lower than the computed maximum neighbouring distance.

In the four data sets, the minimum neighboring distances detected with the Gabriel graphs were 2400 m, 2700 m, 5100 m and 4500 m for the species *A. obstetricans, N. helvetica, M. jurtina* and *A. parallelepipedus*, respectively. In the *A. obstetricans* data set, the spatial scales maximizing the R^2^ between pairs were 3000 m, 2400 m and 3500 m for the Bray-Curtis genetic distance, HGD1 and HGD2, respectively. In the *N. helvetica* data set, the spatial scale maximizing the R^2^ was 2800 m. In the *M. jurtina* data set, the spatial scale maximizing the R^2^ was 5500 m. In the *A. parallelepipedus* data set, the spatial scales maximizing the R^2^ were 6500 m, 18500 m and 4500 m for the Fst genetic distance, HGD1 and HGD2, respectively.

The following figure provides an illustration of the approach for *A. obstetricans*. The inferred minimum neighboring graph is in black (top right). For each measure of genetic distances, the left panel is the plot of R^2^ with increasing spatial scales (maxdist) indicating with a vertical blue axis the optimal spatial scale of analysis (i.e., maximizing R^2^); the right panel represents the corresponding retained neighbouring network (in blue).

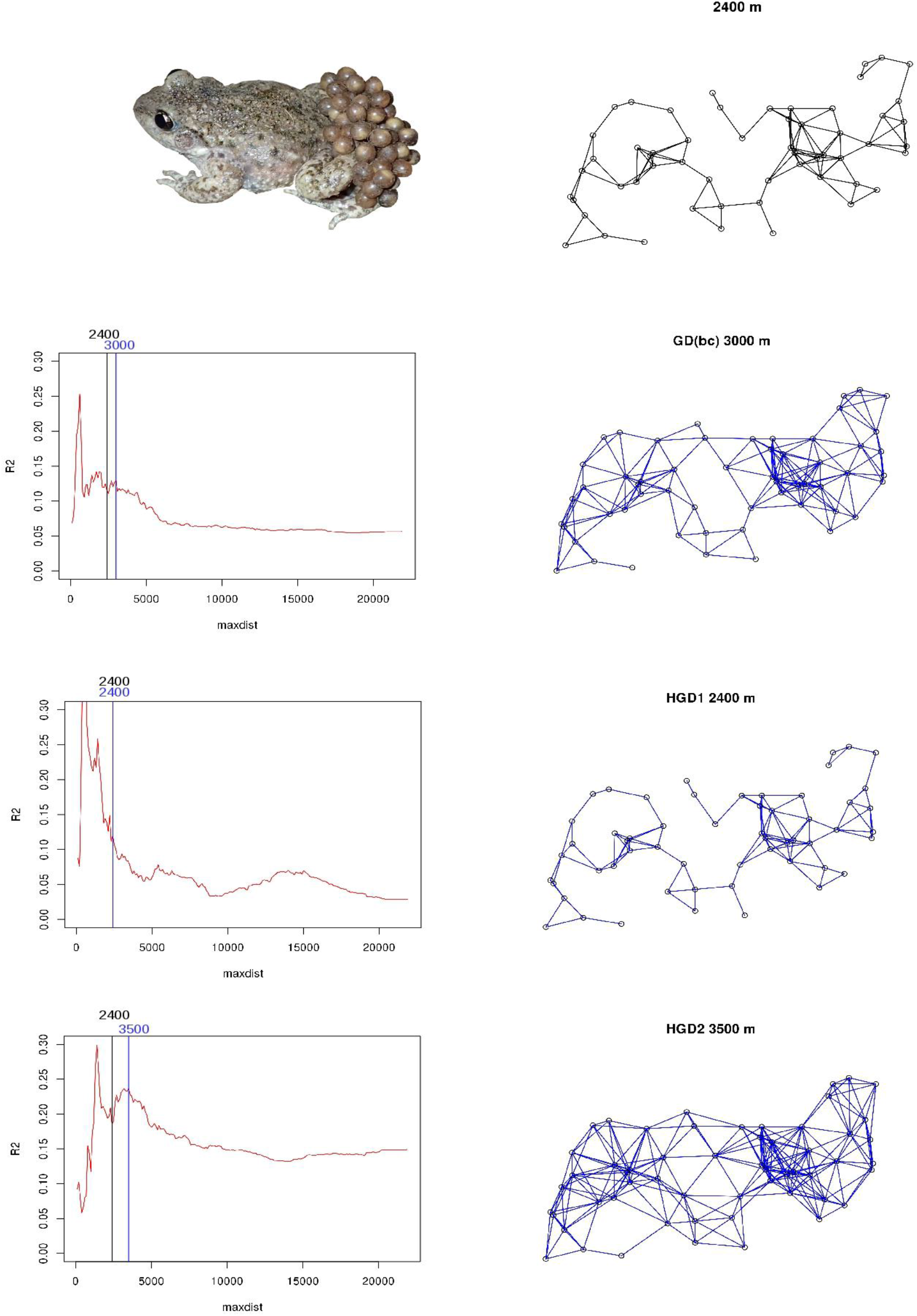

### E. Commonality analyses

In commonality analyses, the effect of each predictor can be decomposed into a unique (U) and common (C; shared with other predictors) effect. For a given predictor, the sum of unique and common effects corresponds to the total contribution (T), equal to its squared zero-order correlation with the dependent variable (U + C = T = r^2^). Therefore, CA represents a good opportunity to assess the reliability of predictors to explain the dependent variable face to collinearity. The magnitude of suppression among predictors is indicated by negative commonalities. Negative commonalities represent the amount of predictive power that would be lost by other predictors if the suppressor variable was not included in the regression model.

Accordingly, we can distinguish three specific types of suppressor (Conger 1974). (i) A classical suppressor corresponds to a predictor whose unique contribution is totally counterbalanced by its negative common contribution (U + C = 0). (ii) A reciprocal suppressor, also described as a partial suppressor, is a predictor with a negative common effect but that does not counterbalance its unique contribution to the variance in the dependent variable (T = U + C > 0). Finally, (iii) a cross-over suppressor is similar to a partial suppressor but with reversal sign. Cross-over suppressors are detected by a sign inversion between the structure coefficients rs and the beta weights (Prunier et al. 2017).

We performed multiple linear regressions and CA using packages ecodist (Goslee and Urban, 2007) and yhat (Nimon et al. 2008) in R 3.3.2 (R Core Team, 2015). To remove classical suppressors, we discarded predictors presenting low univariate squared correlation against the genetic dependent variables (r^2^ lower than 0.1). Low correlated predictors are likely to act as classical suppressors leading to the distortion of regression coefficients (Prunier et al. 2015). When we discarded those non-informative predictors, we ended up with simplified models containing a reduced number of predictors which were likely to explain the variance in the genetic dependent variables. Predictors that were identified as cross-over and reciprocal suppressors were discarded from our model and subsequent models were ran without these suppressors until no more suppressors could reasonably be discarded from the model (that is, we kept reciprocal suppressors showing a non-negligible unique contribution). We also removed predictors with synergistic (S) association with other predictors, which have a unique contribution to the dependent variable equal to zero but presenting synergistic association with other predictors (C > 0).

### F. Intermediate steps of commonality analyses on vectors

To explain the dependent variable based on the Bray-Curtis genetic distance in *A. obstetricans*, the predictors with a squared correlation (r^2^) with the dependent variable higher than 0.1 were IBD, Altitude, Woodlands, Water, Roads, D6089 and Railway. Among these predictors, Water and Railway were cross-over suppressors and were discarded from subsequent analysis. To explain the first level of hierarchical genetic distances (HGD1) in *A. obstetricans*, the predictors with a r^2^ higher than 0.1 were IBD, Woodlands, Water, Crops, Roads and Railway. IBD was a suppressor with synergistic association with other predictors. Water was a cross-over suppressor. These two predictors were discarded and the final model comprised four predictors: Woodlands, Crops, Roads and Railway. To explain the second level of hierarchical genetic distances (HGD2) in *A. obstetricans*, the predictors with a r^2^ higher than 0.1 were IBD, Woodlands, Urban, Roads, D6089 and Motorway. IBD and Urban were cross-over suppressors and were discarded from subsequent analyses.

In the *N. helvetica* data set, only three predictors had a r^2^ higher than 0.1: Roads, Motorway and Railway. There was no suppressor among these three predictors and all were used in the final model.

In *M. jurtina*, five predictors had a r^2^ higher than 0.1: IBD, Woodlands, Grasslands, D6089 and Power line. Grasslands was a cross-over suppressor and the roads D6089 was a partial suppressor. These two predictors were discarded from subsequent analysis resulting in a final model with three predictors: IBD, Woodlands and Power line.

To explain Fst in *A. parallelepipedus*, six predictors had a r^2^ higher than 0.1: Altitude, Grasslands, Water, Urban, Roads and Motorway. Water, Urban, Roads and Motorway were cross-over suppressors. All were discarded from subsequent analysis. Only two predictors remained in the final model: Altitude and Grasslands. To explain the first level of hierarchical genetic distances (HGD1) in *A. parallelepipedus*, we retained the predictors: Grasslands, Water, Crops, Urban, Roads and D6089 (r^2^ > 0.1). Grasslands, Crops and Urban were cross-over suppressors and Water was a suppressor with synergistic association with other predictors. Therefore, we retained only Roads and D6089 to explain the dependent variable in the final data set. To explain the second level of hierarchical genetic distances (HGD2) in *A. parallelepipedus*, we retained the predictors: Altitude, Roads, D6089, Motorway and Gas pipeline (r^2^ > 0.1). The predictor Roads was a suppressor with synergistic association with other predictors and was discarded from subsequent analysis.

The following figures provide the runs of identification of unnecessary predictors for each species and each genetic dependent variable DV (GD: genetic distance either calculated with the Bray-Curtis (bc) dissimilarity index for individual-based method or Fst for population-based method; HGD1 and HGD2 for hierarchical genetic distance based on first and second level of STRUCTURE outputs, respectively). Distance stands for the spatial scale retained in our analyses (Appendix D). Results of the different runs of multiple linear regressions (predictors, structure coefficient rs and standardised coefficient β), in addition to parameters derived from CA: unique (U), common (C) and total (T) contributions of predictors to the variance in the genetic dependent variable. The rationale for withdrawal of predictors (Ra) is the following: CO: cross-over suppression; S: synergistic association with other predictors; PS: partial suppression (or reciprocal suppression). All predictors (IBD: isolation by distance; D6089: a large country road; Urban: urban areas; see Appendix C for additional information on predictors) were coded as resistance. In bold: parameters allowing the identification of unnecessary predictors and suppressors. Note that situations of classical suppression were avoided by discarding any predictor with a squared zero-order correlation < 0.1.

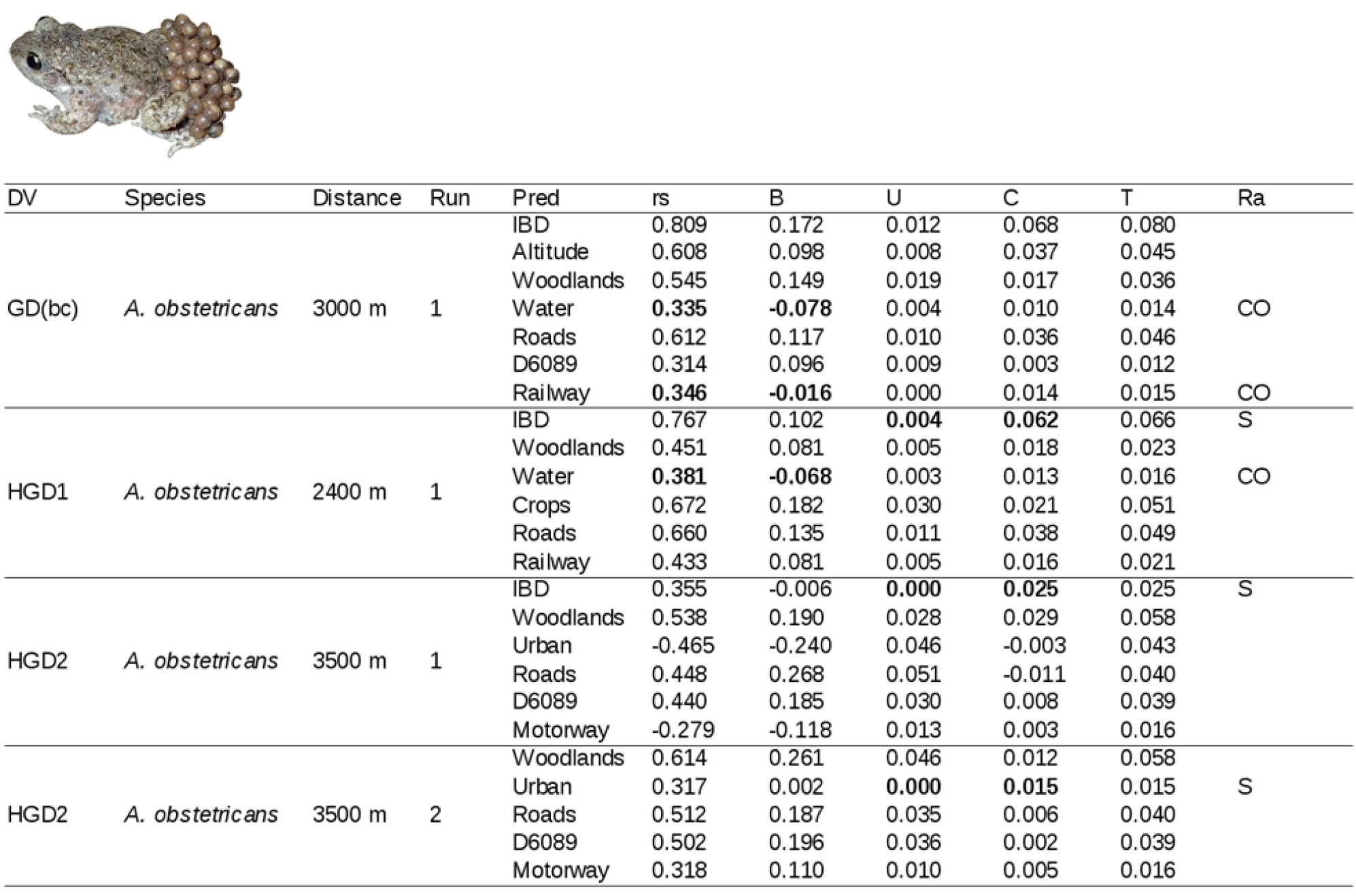
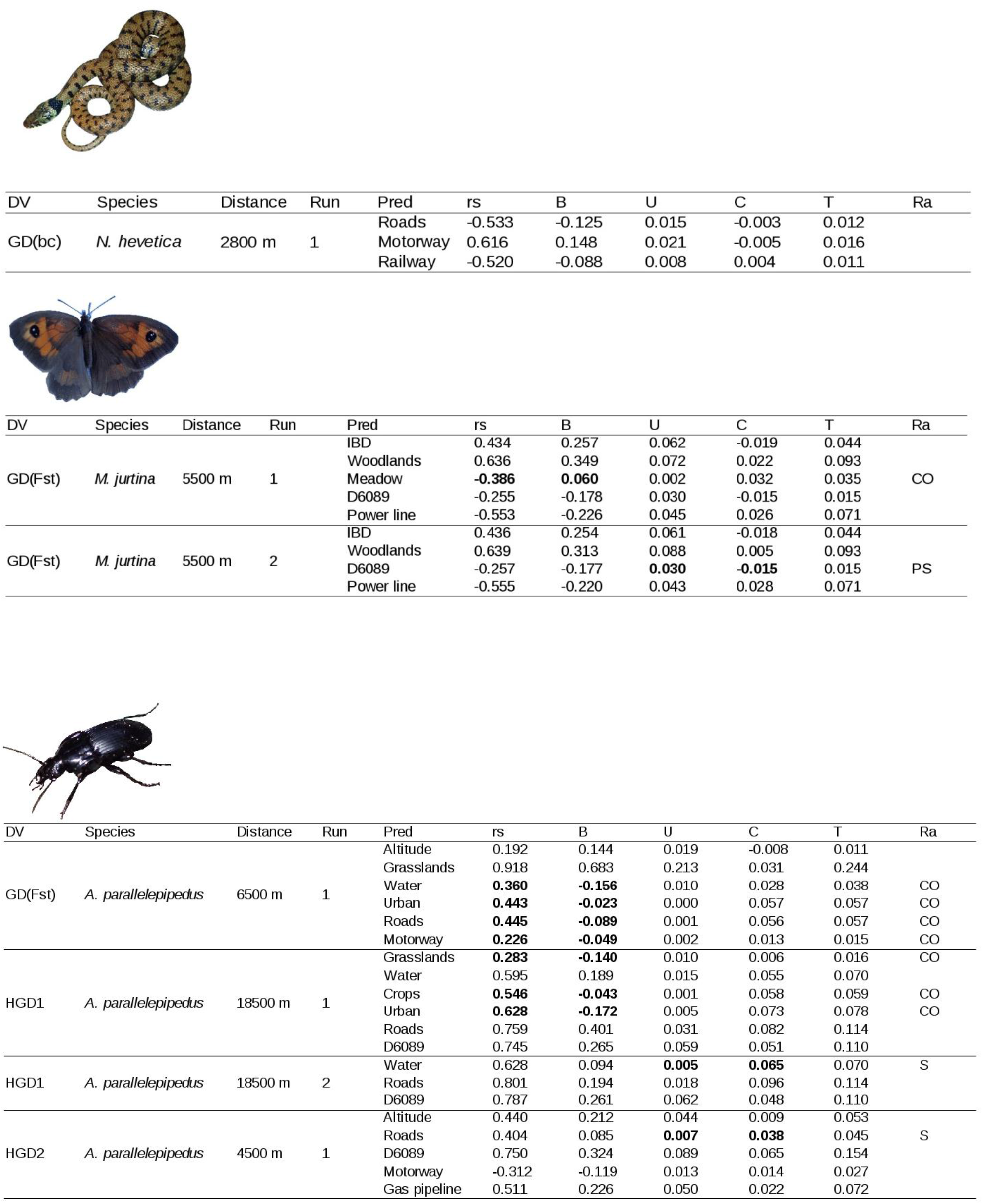

### G. Correlations among final predictors

Matrices of Pearson’s correlation coefficients among final predictors depending on the genetic dependent variables. The genetic dependent variables are genetic distances (GD) based on the Bray-Curtis dissimilarity index (bc), Fst or hierarchical genetic distances at the first and second levels of STRUCTURE outputs (HGD1 and HGD2). The variance inflation factors (VIF) are presented for each predictor.

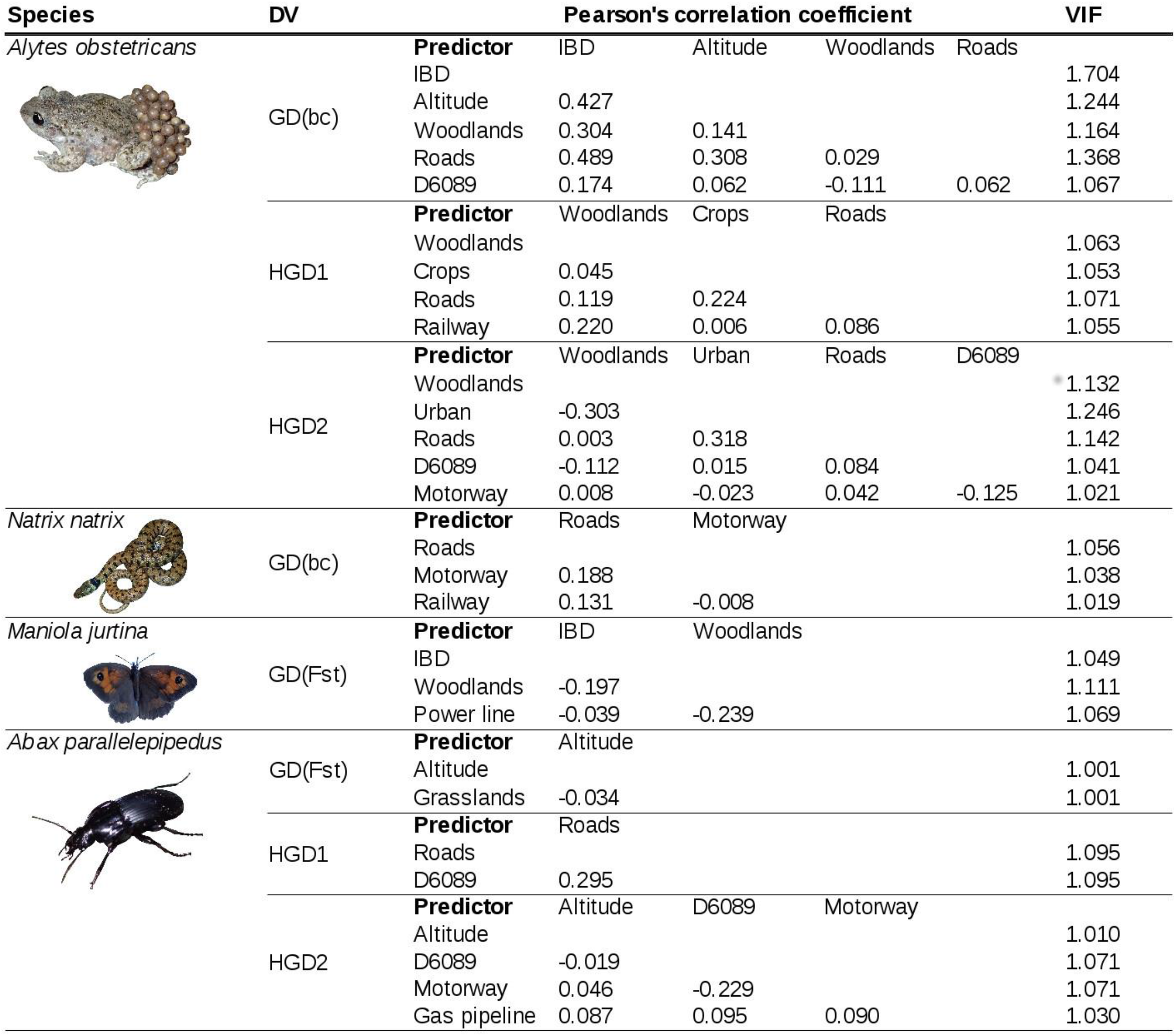

### H. Supplementary results and discussion

#### Alytes obstetricans

With the classical genetic distances, natural predictors (IBD, Altitude and Woodlands) explained most of the dependent variable’s variance (67% of the averaged unique contributions). Woodlands was the landscape element with the highest unique contribution to the genetic distances (U = 0.018).

At the first level of hierarchical genetic distances (HGD1), crops was the predictor with the highest contribution to the dependent variable (U = 0.032) followed by Roads (U = 0.024). In this model, Woodlands was also associated with an increase of genetic distances but was the predictor with the lowest unique contribution (U = 0.010). Railway was associated with an increase of genetic distances with a unique contribution of 0.011 to the dependent variable.

At the second level of hierarchical genetic distances (HGD2), a higher portion of the dependent variable’s variance was explained by our model: 20%. The final model comprised five predictors: Woodlands, Urban, Roads, D6089 and Motorway. Woodlands, Roads and the road D6089 were associated with an increase of genetic distances (positive beta values) but urbanization and the motorway had negative beta values indicating that these two predictors were associated with a reduction of genetic distances. Urbanization was the landscape element affecting the highest part of the dependent variable’s variance (U = 0.047). Woodlands, Roads and the road D6089 were all associated with an increase of genetic distances in this model with unique contribution of 0.031, 0.033 and 0.037, respectively.

#### Maniola jurtina

In this species, woodlands were associated with an increase of genetic distances indicating a barrier effect (positive beta values) and explained most of the variance (U = 0.089). The rest of the explained variance was due to isolation by distance (IBD, U = 0.066). Therefore, the entire variability detected in the butterfly genetic distances was explained by natural predictors.

#### Abax parallelepipedus

With the classical genetic distances, two final predictors explained the dependent variable: Altitude and Grasslands. Altitude did not significantly explain genetic distances (95% confidence intervals included 0). Therefore the variance explained by our model was only due to grasslands associated to an increase of genetic distances indicating a strong barrier effect (U = 0.248).

When using the first level of hierarchical genetic distance (HGD1), the linear regression explained 17% of the dependent variable’s variance. HGD1 was fully explained by predictors associated with an increase of genetic distances in the ground-beetle (positive beta values): the secondary road network (U = 0.063) and the country road D6089 (U = 0.059).

When using the second level of hierarchical genetic distance (HGD2), the linear regression explained 27% of the dependent variable’s variance. Four predictors remained in the final model: the altitude, the road D6089, the motorway and the gas pipeline. The 95% confidence interval around the beta value of the motorway included 0 indicating that the motorway was not significantly explaining HGD2. The three remaining predictors were all associated with an increase of genetic distances (positive beta values). The road D6089 was explaining the highest part of the variability (U = 0.114) suggesting a strong barrier effect of this infrastructure on gene flow. The gas pipeline and Altitude had both a unique contribution to the dependent variable of 0.049.

Infrastructures were not the only landscape elements affecting gene flow in the studied species. Half of the explained genetic variability was, in fact, due to non-LTIs features (Fig. 3A in main text). The non-LTIs features influencing gene flow in *A. obsetricans* were isolation by distance (IBD), altitude differences, crops, woodlands and urban areas (Table 1). Despite classical knowledge on amphibians (VanBuskirk 2012), we revealed that woodlands are a strong driver and is a major barrier to gene flow, affecting classical genetic distances (bc), as well as the first and second hierarchical levels (HGD1 and HGD2). Several hypotheses can be suggested to explain this observation. Individuals may be reluctant to move through woodlands because of inadequate soil characteristics, higher predation level, mitigation of their calling calls due to dense vegetation or absence of optimal breeding water bodies. We were able to detect IBD in this study area that was not detected for the same species in Spain (Garcia-Gonzalez et al. 2012) probably because they used mitochondrial DNA instead of microsatellites which are less variable at small geographical scales (Kohn et al. 2006). Individuals separated by high altitude differences were more genetically distant than individuals sampled at similar altitude level. This result could be linked to a hydrology gradient with individuals sampled in the same water catchment more prone to be close genetically. Crops impeded gene flow at the first hierarchical level (HGD1). A similar result was found for the frog *Rana temporaria* in Germany (Lenhardt et al. 2017). Individuals may be unwilling to cross this landscape feature or be killed while crossing crops because of pesticide exposures (Bruhl et al. 2013) or dehydration risk. Finally, urban areas are landscape elements promoting gene flow in *A. obstetricans*. Urban areas are usually considered as inappropriate habitats, limiting gene flow in amphibians (Goldberg et al. 2010, VanBuskirk 2012). Our result could be explained by the habitat requirements of this species. Old farmhouses are ideal habitats because they combine permanent water bodies (watering trough, cattle ponds, wells, *etc*.), open areas and shelters (stone walls, rubble piles, sand piles, tarps, *etc*.). In the rural landscape studied, old farmhouses are the main urban features with few small villages. It is likely that in more intensive landscapes with large towns, this genetic pattern would differ.

In our study area, the genetic structure of *N. helvetica* was weak. The software STRUCTURE detected only one cluster (interpreted as a single population) indicating that gene flow through this landscape was important. This result may explain the low proportion of the genetic variance explained by landscape features (4% of the variance). In a comparable landscape in Switzerland, (Meister et al. 2010) also found that grass snakes belong to a single population. In this study, we found that *N. helvetica* gene flow was affecting only by infrastructures (roads, motorway A89 and the railway). In seems that, at the local scale, grass snake dispersal is not affected by intensively used landscape features such as crops or urban areas (Wisler et al. 2008, Meister et al. 2010, Meister et al. 2012b). Isolation by distance explains the genetic variance at the regional level (Meister et al. 2012) and genetic structuring can probably only be detected at large spatial scales (Kindler et al. 2013, Pokrant et al. 2016, Kindler et al. 2017, Kindler et al. 2018).

Compared to a previous individual-based study that explained less than 5 % of the genetic variance in three sites across France in the butterfly *M. jurtina* (Villemey et al. 2016), we were able to explain about 20% of the variance when using a population-based method and a restricted spatial scale (maximum neighbouring distance = 5500 m). STRUCTURE was not able to find any genetic structure in the data, probably because of high abundance, low specialization and great dispersal capacity in this butterfly (Villemey et al. 2016). Interestingly, we were able to detect an isolation-by-distance effect. This IBD effect was not detected in (Villemey et al. 2016) with pairwise distances up to 60 km apart. We found that woodlands were impeding gene flow in *M. jurtina*, a result similar to (Villemey et al. 2016). The absence of sunlight and the dense vegetation may limit the movements through woodlands.

Unlike Marcus et al. (2015), we found a strong genetic structure in the ground-beetle *A. parallelepipedus* within the studied landscape. The explained proportion of the classical Fst genetic distance was due to grasslands acting as barrier to gene flow. This result is linked to previous studies showing that this species intentionally avoids open fields such as grasslands (Charrier et al. 1997, Petit et al. 1998). This encourages the maintenance of hedges in agricultural environments to favour landscape connectivity between woodland patches (Charrier et al. 1997, Petit et al. 1998, Fournier and Loreau, 1999). Altitude affected gene flow at the second hierarchical level (HGD2), but its effect was modest (Table 1 in main text). In any case, the fragmentation of woodlands due to land conversion, roads or other kind of LTIs could lead to strong isolation of ground-beetles populations. Population abundance are high in this species (Loreau and Wolf, 1993, Keller et al. 2004) but its dispersal capacity is very limited (Charrier et al. 1997, Brouwers and Newton, 2009). Therefore, populations which are not linked by dispersal may suffer from genetic isolation (Fahrig and Rytwinski, 2009, Beyer et al. 2016).

